# Random subsamples of animal populations can reveal intrinsic differences in sociality with key implications in ecology, conservation and disease transmission

**DOI:** 10.1101/2024.11.07.622426

**Authors:** Kimberly Conteddu, Prabhleen Kaur, Michael Brown, Julian Fennessy, Stephanie Fennessy, Emma Hart, Bawan Amin, Anna David, Laura L. Griffin, Jane Faull, Stefano Grignolio, Francesca Brivio, Amy Haigh, Liam Kirwan, Michael Salter-Townshend, Simone Ciuti

**Affiliations:** Laboratory of Wildlife Ecology and Behaviour, University College Dublin, Dublin, Ireland; School of Biology and Environmental Science, University of Dublin, Dublin, Ireland; School of Mathematics and Statistics, University College Dublin, Dublin, Ireland; Giraffe Conservation Foundation, P.O. Box 86099, Eros, Windhoek, Namibia; Faculty of Social and Behavioural Sciences, Utrecht University, Utrecht, Netherlands; University of British Columbia, Department of Forest Resources Management, 2424 Main Mall, Vancouver, V6T 1Z4, Canada; Department of Life Science and Biotechnology, University of Ferrara, via Borsari 46, Ferrara, 44121, Italy; Department of Veterinary Medicine, University of Sassari, via Vienna 2, Sassari 07100, Italy

**Keywords:** social network analysis, wildlife, subsampling, conservation, zoonoses, animal behaviour, animal personality, climate change

## Abstract

Animal populations are under mounting stress from the dual threats of climate change and rapid global human population growth, raising significant concerns about declining wildlife and the rising risk of zoonotic diseases. In many species, social interactions can be a highly plastic suite of behaviours that are responsive to these disturbances and are consequential to other processes like disease transmission and population dynamics. Studying social interactions can be challenging in that researchers often rely on wildlife population subsamples due to practical constraints and costs, which can introduce biases in the reliability of social network metrics. We investigated the extent to which subsamples can depict intrinsic characteristics of wildlife populations using data from three distinct species: peri-urban fallow deer, Alpine ibex and Angolan giraffe. We showed that random subsamples of these populations could still reveal differences in their social behaviour, indicating that, as long as researchers have a reliable estimate of population size, subsampling animal populations can be an effective and precise method to infer their sociality and offer valuable empirical data for management, conservation and zoonotic disease ecology. Furthermore, we demonstrate that non-random sampling, influenced for instance by animal personality and related trappability, can introduce significant biases in social network estimates. These findings underscore the importance of accounting for sampling biases in social network analysis and offer a robust framework for using partial networks in ecological studies and conservation management.

## Introduction

Global warming is quickly changing the world’s climate, causing extreme weather and resulting in permanently altered ecosystems. Temperature changes have been shown to disrupt wildlife populations [1] by changing migration and breeding patterns [2, 3] as well as habitat suitability [4]. For example, rising temperatures can decrease the availability of suitable refuges, forcing animals to increasingly share refuges and potentially alter parasite and disease transmission [5,6]. Additionally, habitat loss can disrupt disease transmission by forcing wildlife to move closer to semi-urban and urban areas, increasing interactions with humans and domestic animals (e.g., caused by wildlife feeding and improper waste disposal) [7, 8]. Zoonotic disease transmission poses significant health risks to humans and welfare concerns for both domestic and wildlife species [9, 10], with substantial economic consequences, as demonstrated during the COVID-19 pandemic [11, 12]. As such, understanding and predicting the emergence of zoonotic diseases in the Anthropocene has become a top priority to safeguard public and animal health worldwide.

Behavioural changes in wildlife populations under human pressures can critically alter zoonotic disease transmission by increasing contacts between wildlife species, domestic animals and humans [13]. Social Network Analysis (SNA) is a powerful tool to study spatial and temporal changes in behaviour between wild animals [14], helping us understand how they might be altered by human perturbations (i.e., with both direct - e.g., habitat loss, human-wildlife interactions - and indirect effects - e.g., global warming, [4, 15]). Animal social networks can be built using direct observations (e.g., focal observations), where all animals are observed and data on interactions between animals are collected by an experienced observer [16]. However, this can be time-consuming and it is usually not possible to collect high-frequency data (i.e., usually such data collection types are only feasible to conduct once a week/month when investigating long-term spatial and temporal trends [16]). Therefore, equipment such as GPS trackers can be useful for interaction analysis, since they can track multiple animals continuously and simultaneously, allowing the collection of extremely detailed animal interaction data (i.e., data collected up to 1 minute or less) [17]. However, this data collection type presents two main problems. First, placing GPS trackers on wild animals can be labour intensive and expensive (i.e., other concerns include wildlife welfare, battery life, problems capturing animals in inaccessible areas and difficulty in attachment due to wildlife size and morphology, [18]), therefore usually only a small percentage of the population can be tracked at any one time [17]. This can engender significant biases when building social networks, since many nodes (which represent individuals) are missed and could present a completely misleading ecological picture, making the comparison of social networks challenging (e.g., looking at before-and-after scenarios such as human perturbation events or conservation management implementations) [19, 20]. Another problem when collecting data using GPS trackers is capture bias (i.e., animal trappability). When animals are captured, there is often a sampling bias towards certain personality types over others [21–25], which can be characterised by different sociality and affect the structure of social networks [26], thus giving a false representation of the true relationships between individuals of a focal population.

Hence, it is crucial to consider the uncertainties involved in using subsamples of the population to construct animal social networks (i.e., partial networks) and to evaluate whether these partial networks can effectively inform decision-making instead of relying on full population data. To date, there has been limited research focusing on using partial networks to make inferences about entire populations [19, 20]. Previous research has mainly focused on estimating the uncertainty surrounding single partial networks. One such example is a study by Silk *et al.* [19] which investigates the effect of subsampling on social network metrics through simulations. In addition, Kaur *et al.* [27] used real data to determine the effect of subsampling in wildlife populations and implemented a protocol to account for partial network uncertainty. However, research to date has not investigated the use of subsamples in comparing animal social networks to detect sociality changes in a wildlife population (under human perturbations and climate change pressures, for instance) nor the effect of capture bias on social network metric estimates. Therefore, the first goal of this study was to investigate whether differences between networks can be captured by only monitoring a subsample of individuals in a population. By randomly reducing the number of animals monitored, we aimed to understand how differences in social behaviour are captured using real interaction data and to identify the minimum subsample percentage needed to detect differences between networks that accurately represent the entire population. The second goal was to examine the impact of capture bias on social network metrics by comparing networks where animals are selected randomly to those where selection is biased towards a specific personality type and to determine if this bias persists regardless of subsample size.

To achieve our first goal, we used observation data from three anthropogenically-disturbed ungulate populations occupying different ecological niches: a fallow deer (*Dama dama*) population living in Phoenix Park, Dublin which is experiencing unprecedented human pressure from feeding by park visitors; a population of Alpine ibex (*Capra ibex*) living in the Italian Alps, particularly affected by raising summer temperatures; and, an Angolan giraffe (*Giraffa giraffa angolensis*) population living in the Namibian desert, an extreme environment which is predicted to become more challenging under future global warming scenarios, leading most likely to aridification [28]. For each population, a significant portion of the entire population was tracked (varying from 30% to 85%). For each species and related ecological context, we asked three ecological questions, tested for differences in the social networks for the entire population and verified whether random subsamples could retain the intrinsic differences of the whole population. Specifically:

- For the fallow deer population, we looked at the effect of management on the social network of fallow deer. Between 2018 and 2019 the park implemented a management plan to decrease the feeding of deer by visitors [29]. Since visitors have been seen significantly disrupting the behaviour of deer in the park (i.e., artificial feeding and related disruption to social behaviour), we were interested in the effect on the relationships between deer and their sociality. We expected deer to be less aggregated in 2018 before the management implementation (i.e., higher levels of disruptions by humans continuously disrupting groups, [29]) compared to 2019, after the management implementation (i.e., lower levels of disruptions by humans allowing deer to form and maintain larger groups, [29]). We, therefore, tested for social network differences between 2018 and 2019 in the entire population as well as in random subsamples;
- For the Alpine ibex population, a cold-adapted species affected by high temperatures [30], we aimed to disentangle the effect of hotter conditions on ibex sociality, and whether population subsamples could retain such differences. We expected ibex to be more clustered together at lower temperatures (i.e., when they can gregariously graze in Alpine pastures at low altitude and of high quality) compared to higher temperatures (i.e., when ibex are forced by thermal stress to abandon high-quality feeding clusters and to use sub-optimal area);
- For the Angolan giraffe population, seasonality and daily variations in temperature and food availability significantly affect their sociality [31–34]. Therefore, we aimed to compare network changes depending on the season (when temperatures and related water and food availability can change significantly in such a hyper-arid environment) and time of day. We expected the season to influence aggregation depending on food availability (i.e., higher clustering with lower food and water availability). Furthermore, because giraffe tend to form large feeding clusters in the morning as opposed to smaller and more bonded units in the evening [31], we expected giraffe connections to be tighter in the evening compared to the rest of the day. Again, as for the other focal populations, we tested for these differences in the full network and investigated whether random subsamples could still depict the intrinsic characteristics of giraffe social networks.

To achieve our second goal, we used the Phoenix Park fallow deer population since we could avail of behavioural data on feeding interactions with humans for the entire population. Previous studies in the park showed that only ∼20% of the population consistently approach closely and accept food from humans (bold individuals), even directly from their hands [35]. The rest of the population either occasionally accepts food or actively avoids humans (shyer individuals). Approachable individuals (food acceptors) would be easily immobilised by researchers (e.g., using dart guns or nets) compared to those actively avoiding humans. We used this dataset to simulate a scenario where bolder individuals are more easily trappable, and compared social network metrics between random sampling (i.e., all animals having the same probability of being selected) and biased sampling (i.e., bold individuals being more likely to be selected than random).

## Materials and Methods

All statistical analyses were done using R version 4.4.0 [36].

### Effect of subsampling in comparing wildlife social network metrics

We first sought to understand how metrics behave when we only have a subsample of the wildlife population and if they can still be used to detect differences in social networks (e.g., seasonal/temporal variation, human perturbations etc.). We used empirical data from three wild wildlife populations with extremely different ecological characteristics: fallow deer, Alpine ibex and Angolan giraffe (see Appendix S1 for a detailed explanation of the three datasets and their data collection). All populations have been regularly surveyed every month across multiple years via direct observations of individually recognisable individuals (ear-tagged deer and ibex; individual coat patterns in giraffe) [30, 32, 35]. The overall population estimates in each case were available during the study period (See Appendix S1), meaning that we had a precise idea of the proportion of the population monitored (up to 86% in ibex, Appendix S1). Given this accurate picture of all three populations, our random subsampling exercise described below emulates researchers randomly selecting and tagging a small portion of a population (e.g., 10% of the individuals to be ear-tagged or even fitted with GPS collars). Since we had the full network, we could then assess whether researchers would be able to make any meaningful inference on their target population by using their limited samples and whether these subsamples could provide a meaningful estimate of population-level network metrics.

#### • Fallow deer - wildlife feeding

In the fallow deer population we tested for differences in sociality before and after the introduction of management controls on wildlife feeding (May 2018 - December 2018: before management controls; January 2019 - December 2019: after management controls [29]). Since the number of observations and groups observed in 2019 was greater than in 2018 (n. observations: 19,768, n. groups: 445) we conducted a down-sampling for the 2019 dataset where we randomly selected the same number of groups as 2018 (n. groups: 364). We used a total of 14,446 and 15,755 observations to build the 2018 and 2019 social networks, respectively. The 2018 network had 550 nodes while the 2019 network had 556 nodes. We matched the number of nodes by randomly selecting 550 nodes from the 2019 network to have comparable networks for the statistical analysis described in full below.

#### • Ibex - temperature

The social interactions in the ibex population were expected to be greatly affected by temperature fluctuations, particularly when higher temperatures may disrupt feeding behaviour and gregariousness. We therefore tested how social interactions change between low temperatures (i.e., temperatures below 10°C, the first quartile in the dataset) and high temperatures (i.e., temperatures above 14°C, the third quartile in the dataset). The two ibex datasets created for low and high temperatures had a similar number of groups and observations, therefore no down-sampling was required (n. groups: high (139), low (144); n. observations: high (834), low (974)). The networks created from the datasets had 53 and 51 nodes for high and low temperature respectively, therefore we matched the number of nodes by randomly selecting 51 nodes for the high temperature network.

#### • Angolan giraffe - season and time of day

In the Angolan giraffe population, we tested for seasonal and daily differences in social contacts between individuals as a result of varying temperatures and related water and food availability. Resource availability, as well as sociosexual behaviour, differs between the seasons (i.e., cold-dry season, hot-dry season, wet season) which consequently affects the social interactions between giraffe [31]. In addition, interactions between giraffe change depending on the time of day (TOD, i.e., morning, midday, evening;) likely due to behaviours that aid in body thermoregulation [31]. The dataset from the giraffe data collection included unequal number of groups and observations during the different seasons (n. groups: cold-dry (90), hot-dry (315), wet (107); n. observations: cold-dry (334), hot-dry (1150), wet (347)) and TODs (n. groups: morning (114), midday (336), evening (62); n. observations: morning (430), midday (1207), evening (194)). We decided, therefore, to down-sample the dataset to match the lowest number of seasonal groups (n. groups: 90) and TOD (n. groups: 62). The number of observations, after down-sampling, used to build the social networks for the three different seasons were 334, 344 and 256 for cold-dry, hot-dry and wet season, respectively. As for TOD, we used 242, 236 and 194 observations for morning, midday and evening, respectively. The networks created for season had 129 (cold-dry), 130 (hot-dry) and 123 nodes, whereas for time of day they had 110 (midday, evening) and 104 (morning) nodes. We then matched the number of nodes (as done above for the other populations) to 123 for all the season networks and 104 for all the TOD networks.

#### • Subsampling

The first step of the subsampling analysis was to calculate the weights which would be used to build the wildlife networks. The weights we used indicate the strength of association between an individual and all the other individuals in the population of interest (e.g., a fallow deer and all the other fallow deer individuals in the population). We used the Half-Weight Index (HWI) to calculate the weights for all three wildlife populations (Equation 1). We decided to use this index instead of the simple ratio index following the recommendation of Farine and Whitehead [37, pp. 1149]. They indicate in their study that the HWI index is better suited when “individuals are frequently missed (when they should have been observed)” since it “can provide a less biased estimate of the real rate of association”, which was the case in all the three of the populations used in this study.

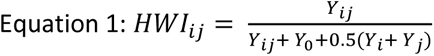

where *Y*_*ij*_ indicates the number of sampling times where animal *i* and *j* were seen together, *Y*_*i*_ indicates the number of sampling times where animal *i* was seen but animal *j* was not, *Y*_*j*_ indicates the number of sampling times where animal *j* was seen but animal *i* was not, and *Y*_0_ the number of sampling times where animal *i* and *j* were seen but in different groups.

We then created all networks (Deer: Management effect networks; ibex: Temperature networks; Giraffe: Season and Time of Day networks) for the three species using the package *<u>igraph</u>* [38]. Once the networks were created, we used the package *<u>aniSNA</u>* [20] to randomly sample nodes from the observed networks (e.g., low and high temperature networks of the ibex population). For each network, we drew 100 random samples, without replacement, of *m* nodes from 95% to 5% of the total number of nodes from the observed network, *N* [20]. Note that the 5% subsampling percentage was not used for the ibex networks since it was too small of a percentage to create a network subsample for this population (N = 51). For each of the 100 samples of all the subsampling percentages, we calculated three metrics: mean degree, mean strength and transitivity (these metrics were chosen since they are reliable even at low sample sizes, see Kaur *et al.* [20]). Since mean degree and strength are directly affected by the number of nodes in the network [20], we decided to scale both metrics by the latter to be able to easily compare these metrics between samples from different subsampling percentages (a full description of the social network metrics computed in this study can be found in Table 1).

**Table 1:** Description of the social network metrics calculated for the three ungulate populations (i.e., fallow deer, Alpine ibex, Angolan giraffe) examined in the study.

| Statistic | Description |
| --- | --- |
| Scaled Mean Degree | Mean degree measures the average number of individuals all animals have interacted with. In this case it is also scaled by the total number of individuals (nodes) in the network. A higher number indicates that on average the individuals of the population interact with more animals, while a lower number indicates that on average the individuals of the population interact with fewer animals. |
| Scaled Mean Strength (Weighted) | Mean strength measures the average strength of association between all individuals in the population. The strength in this case is also weighted (weights are assigned to all associations instead of being just presence and absence of association between individuals, the weights in this study are calculated using the HWI index) and scaled (by the total number of individuals, nodes, in the network). A higher number means that the population is more socially connected, while a lower number indicates that the population is less socially connected. |
| Transitivity | Measures how clustered the wildlife population is. It is calculated by counting the number of complete triangles (three individuals interacting with each other) in relation to all possible triangles. A higher number means that there are more groups in the population that are highly socially connected, a lower number means that there are fewer groups in the population that are highly socially connected. |

Once we calculated metric values (scaled mean degree, scaled mean strength, transitivity) for all the network subsamples, we used a Welch Two sample t-test to compare the network metrics between the deer networks (two networks: year 2018 *vs* year 2019) and ibex networks (two networks: >14 °C *vs* < 10 °C) and a linear model to compare the season (three seasonal networks: hot-dry, cold-dry, wet) and TOD networks (three daily networks: morning, midday, evening) of the giraffe population. For the deer and ibex networks, we used the function *t.test* from the R package *<u>stats</u>* [36]. For the giraffe population networks, we used the function *lm* from the package *<u>stats</u>* [36].

The statistical tests mentioned above allowed us to test whether there was still a significant difference between social network metrics when using population subsamples. A subsample, however, may allow us to detect the difference between two social networks (e.g., a significant difference in the ibex network when temperatures were cooler *vs* hotter), though the social network estimates of the subsample can deviate from the estimates for the full population. Therefore, we investigated whether the metric estimates of network subsamples were still representative of the observed network values from the full population. To do so, we used a linear model (using the *lm* from the package *<u>stats</u>, Equation 2*, and the function *allEffects* from the package *<u>effects</u>* to plot the results, [39]) to study the variation of network metrics along with conditional confidence intervals as a function of subsample dimensions, and we repeated this for all wildlife populations and network types.

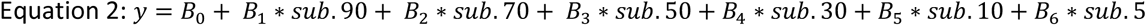

where B_0_ indicates the intercept (the 95% subsampling percentage), B_1_-B_6_ are the estimates for each subsample size indicating the difference in estimates at each subsample size compared to the 95% estimate), sub.90 to sub.5 indicates the subsample sizes or predictors. Note that sub.5 was not included for the ibex networks due to the small sample size.

We then used these models to estimate the minimum population subsample (proportion of monitored individuals) which were able to provide social network estimates not significantly different to those revealed by the entire network. Note that in all statistical analyses, the network metrics corresponding to the baseline full population were computed on 95% of the individuals to allow confidence interval computation.

### Effect of capture bias on social network metrics

We then sought to understand the effect of capture bias on network metrics. We investigated the effect of an increased likelihood of capturing animals with certain behavioural types on network metrics and whether it would be significantly different from random sampling. In addition, we aimed to understand whether this difference was maintained at every subsample percentage.

We used data from the Phoenix Park fallow deer population, which contains behavioural data linked to feeding interactions between deer and people. The deer were previously classified into three behavioural types: consistent beggars, occasional beggars and rare beggars (See Griffin, Haight, Amin, *et al.* [35], for more details on the data collection and analysis conducted to categorise animals into behavioural types). Consistent beggars are animals that consistently approach and accept food offered by people even directly by hand, whereas occasional beggars and rare beggars are animals that only occasionally accept food from people and actively avoid people respectively. For the analysis, we divided the three behaviours into two main categories, where consistent beggars are referred to as bold individuals since they consistently interact with humans, while occasional beggars and rare beggars are referred to as shy individuals since they only occasionally interact with humans or actively avoid them.

We used a total of 85,280 observations and 3,233 groups collected between May 2018 and January 2023 to build the overall social network of deer in Phoenix Park. The network had 751 nodes with 158 nodes being deer that consistently accept food from humans (hereinafter bold deer) and 593 of the nodes being deer that only occasionally accept food or avoid human interactions (a.k.a. shy deer). We calculated the weights in the network, representing the strength of interactions between animals, using the HWI index defined above. We then built a weighted network using the package *<u>igraph</u>* [38]. From the network, we drew 100 random samples, without replacement, of *m* nodes from 95% to 5% of the total number of nodes (N = 751) [20]. The random probability of selecting a node depending on the behavioural types was P = 0.21 for bold individuals and P = 0.79 for shy individuals. We then wanted to compare this random sampling to a capture biased sampling where bold individuals are more likely to be picked compared to shy individuals. Therefore, given that it is common for park visitors to approach and touch bold deer (see Griffin, Haight, Amin *et al.* [35]), we arbitrarily biased the sampling by increasing the probability of sampling bold individuals to P = 0.91 while shy individuals had a probability of being selected of just P = 0.09. We drew 100 biased samples, with replacement, of *m* nodes from 95% to 5% of the total number of nodes (N = 751) (see Appendix S2 for additional analysis on the effect of sampling with replacement and without replacement). For each of the 100 samples of all the subsampling percentages for both the random and the biased sampling we calculated the same three metrics as introduced earlier: scaled mean degree, scaled mean strength (weighted) and transitivity. We then used the Paired t-test using the function *t.test* from the R package *<u>stats</u>* [36] to compare the metrics’ means in function of the sampling type (random *vs* biased), to see if the difference between the sampling types remained consistent regardless of the subsampling percentage.

## Results

We described in full detail below the network analysis for the three species (summarised in Figure 1- 2) as well as how subsamples can still detect differences recorded at the whole network level (Figure 3-4), followed by an analysis of how network metrics estimated for subsamples can mimic those recorded for the entire population (Figure 5-6). Finally, we assessed how random sampling vs sampling of more easily trappable individuals has the potential to bias social network estimates in natural populations (Figure 7).

**Figure 1:**
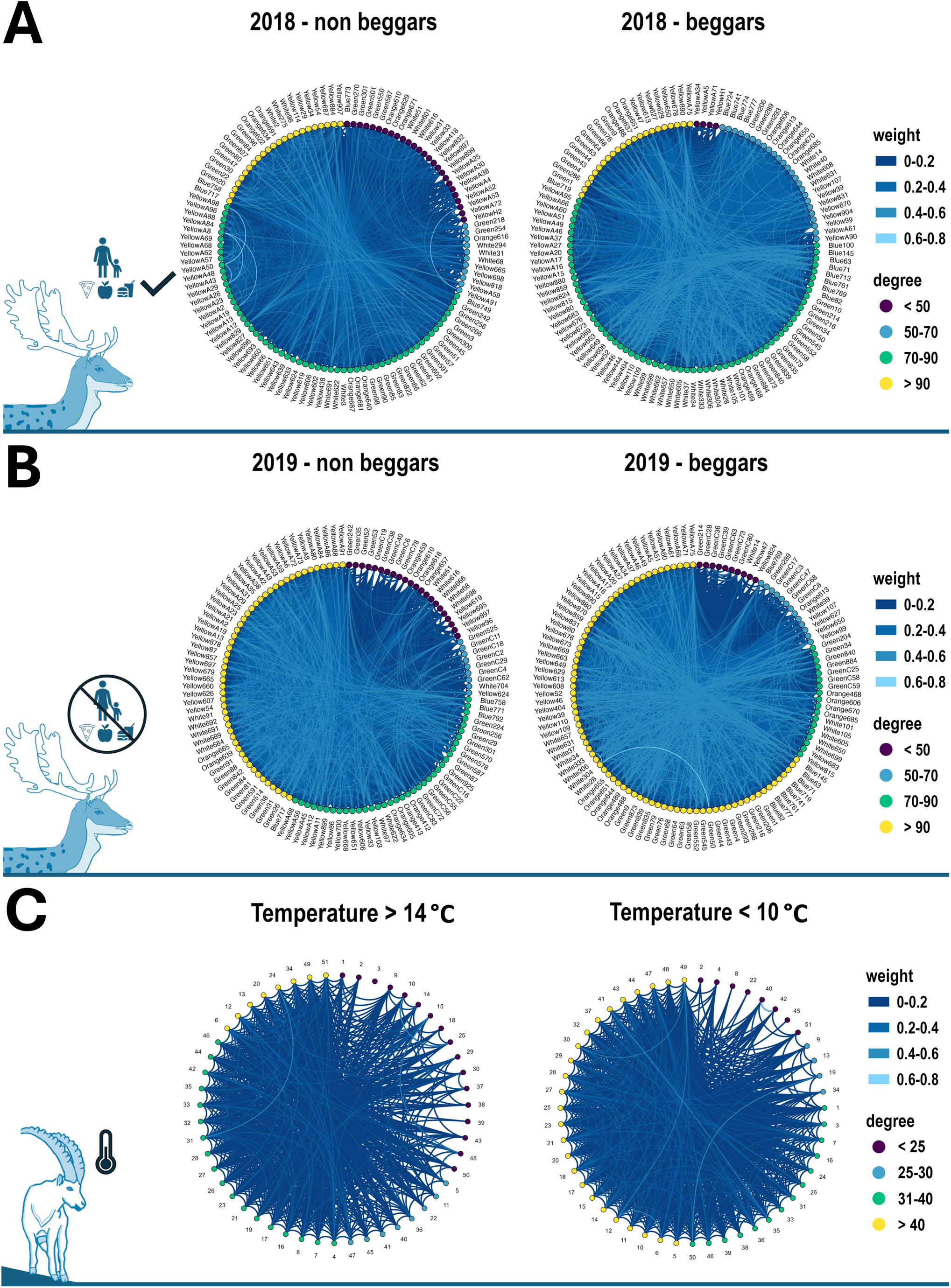
Social networks representing interactions between deer of Phoenix Park, Ireland **A)** before and **B)** after the management implementation aimed to reduce human-deer interactions for both non-beggar/shy individuals - i.e., only occasionally or never beg for food from humans - and beggars/bold - i.e., consistently beg for food from humans; and **C)** social network of Alpine ibex at low - i.e., < 10 °C - and high - i.e., > 14 °C - temperatures. Weight colours indicate the strength of the connections between two individuals, while node colours indicate how many individuals that animal has been in contact with. Note that for the fallow deer networks the number of nodes differed between networks therefore we used the lowest number of nodes (n. nodes = 119, which was the number of beggar individuals for 2018) and randomly selected the same number of individuals for the other networks, so that they would be visually comparable. All species drawings were made by Kimberly Conteddu.

**Figure 2:**
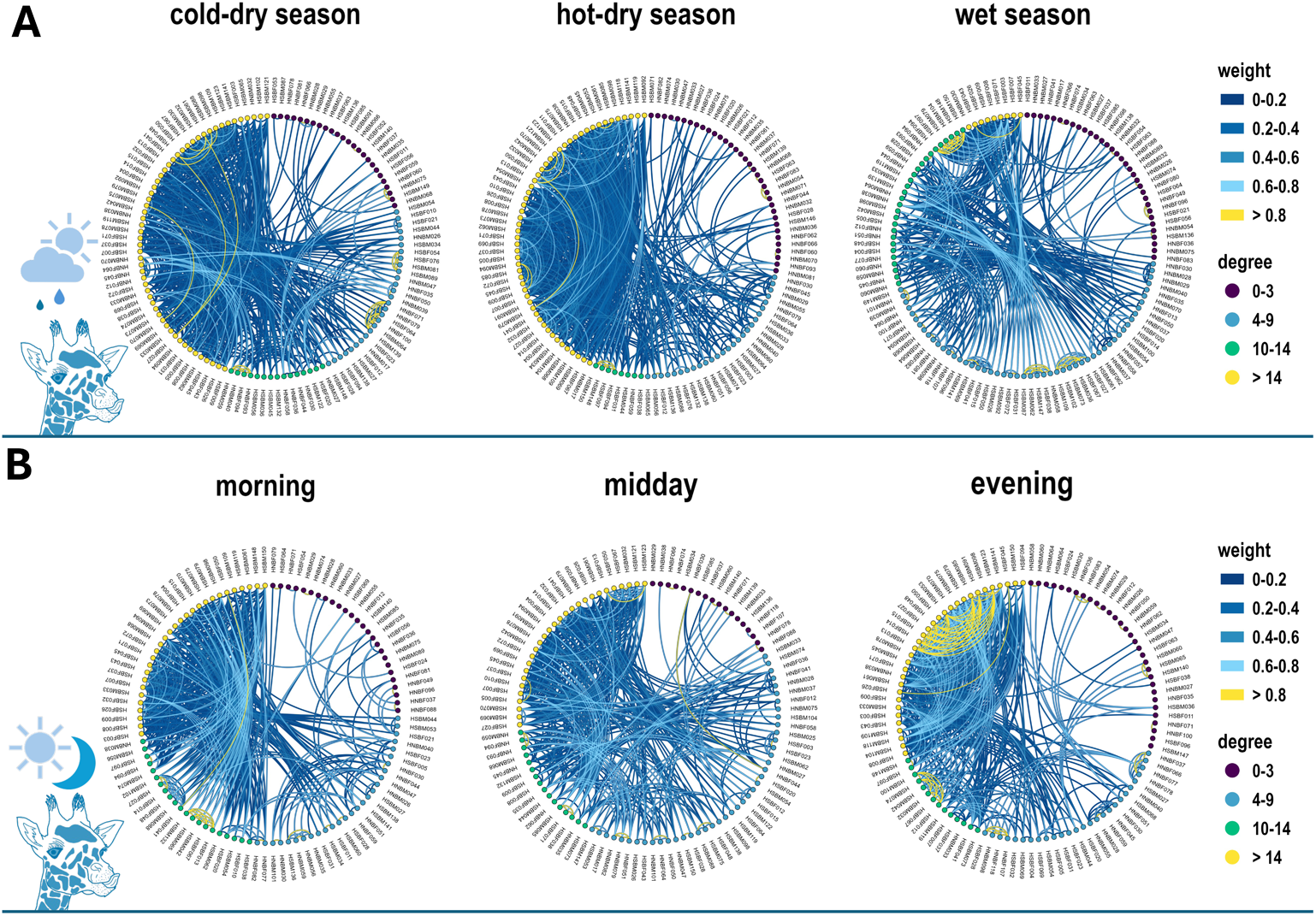
Social networks representing the social interactions between **A)** Angolan giraffe in the Namib desert during the hot-dry, cold-dry and wet seasons and **B)** during different times of day: morning, midday and evening. Weight colours indicate the strength of the connections between two individuals, while node colours indicate how many individuals that animal has been in contact with. All species drawings were made by Kimberly Conteddu.

**Figure 3:**
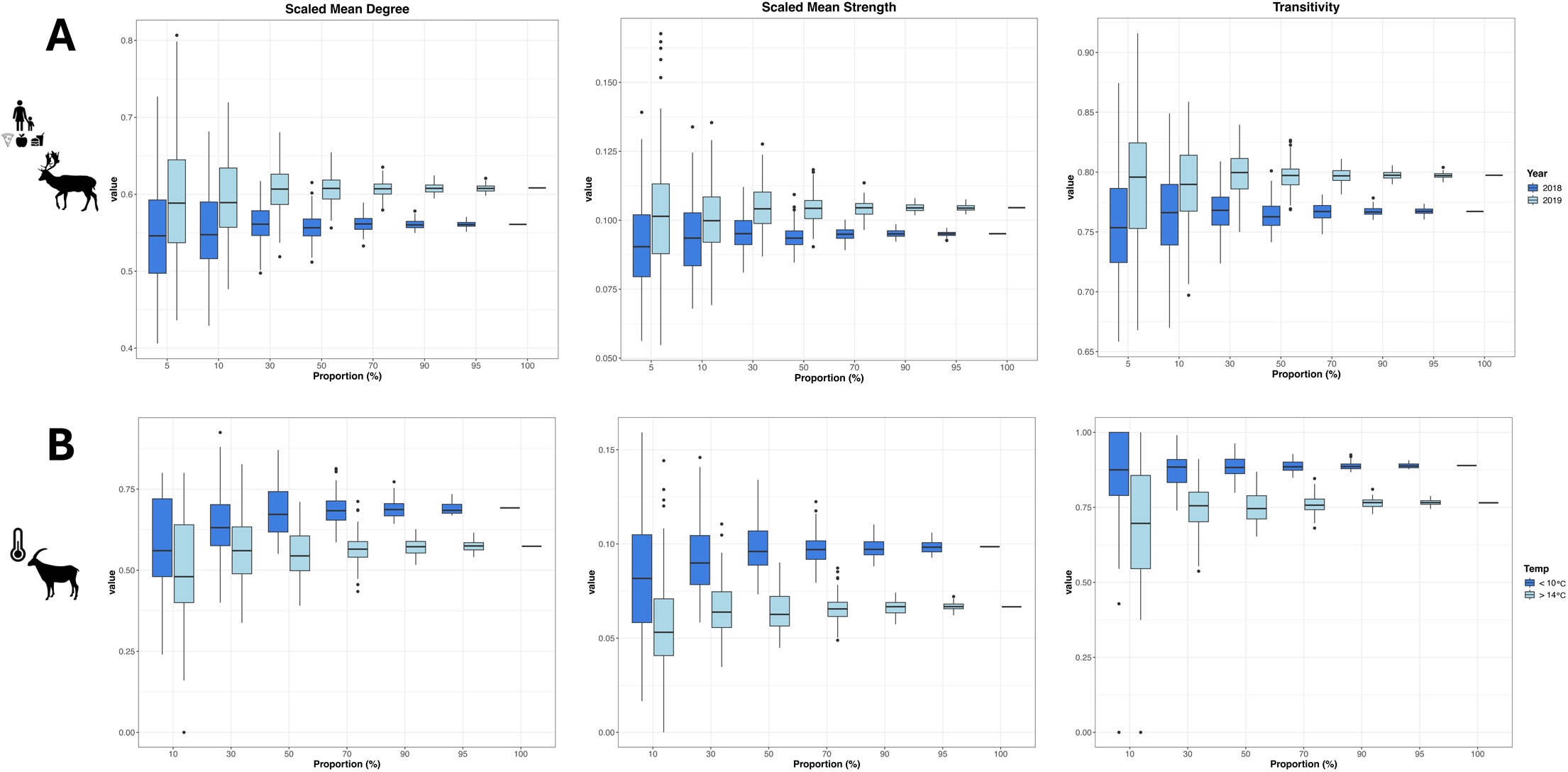
Effect of subsampling on the social networks of **A)** deer before (dark blue; i.e., 2018, higher human disturbance levels) and after the management implementation (light blue; i.e., 2019, lower human disturbance levels); **B)** ibex at low (dark blue; i.e., < 10 °C) and high temperatures (light blue; i.e., > 14 °C). Each boxplot for the three network metrics - i.e., scaled mean degree, scaled mean strength and transitivity - indicates the 95% confidence interval of the network metrics calculated for 100 random samples of the overall population. In addition, for each population, the real network metrics estimates of the overall population networks are shown on the left of each of the plots - i.e., proportion equal to 100%. Animal silhouettes were downloaded from PhyloPic (https://www.phylopic.org/). Deer silhouette is by Anthony Caravaggi under: Attribution-NonCommercial-ShareAlike 3.0 Unported (CC BY-NC-SA 3.0), link: https://creativecommons.org/licenses/by-nc-sa/3.0/. Ibex silhouette is by Steven Traver under: CC0 1.0 Universal (CC0 1.0) Public Domain Dedication.

**Figure 4:**
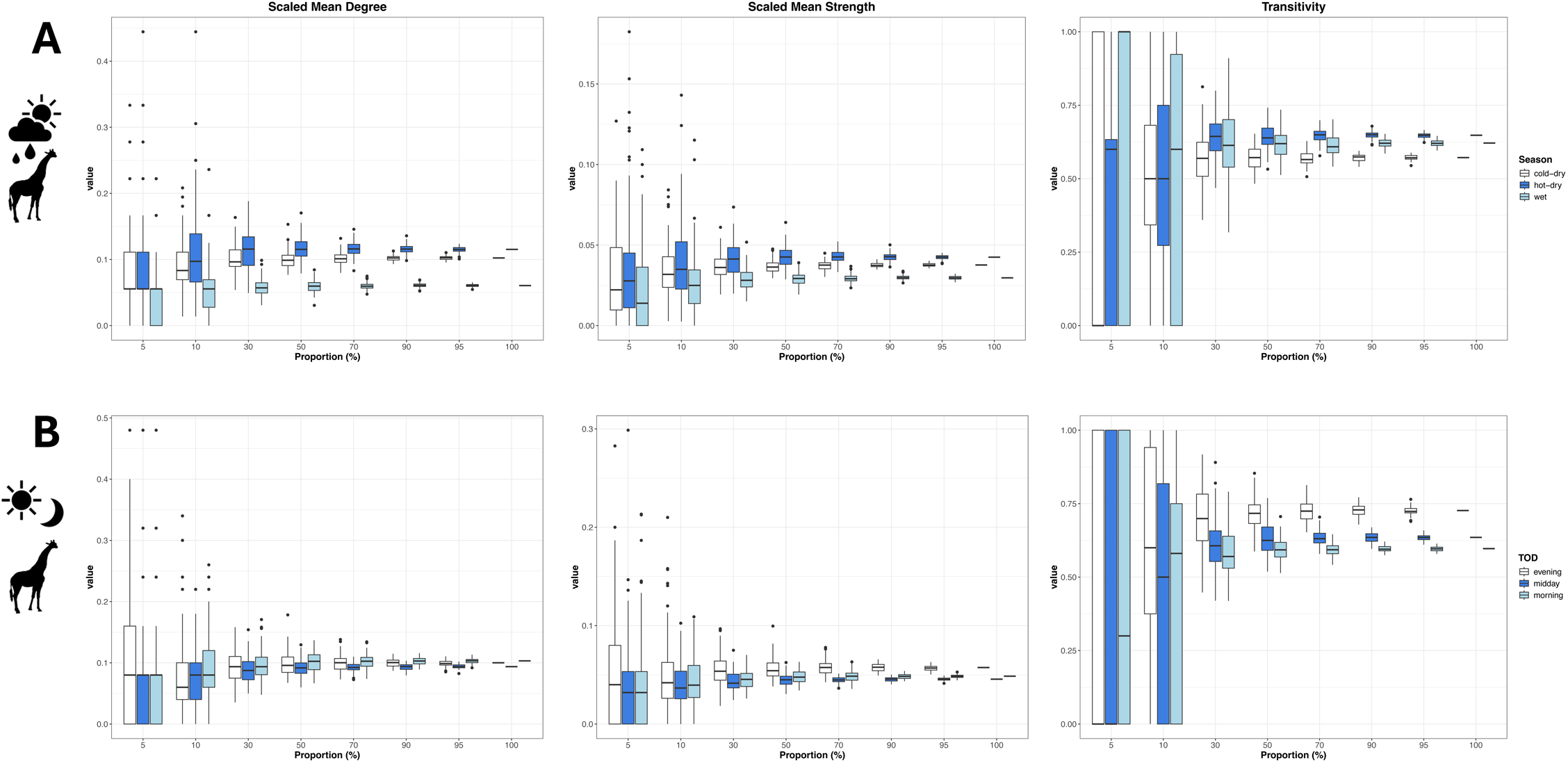
Effect of subsampling on the social networks of **A)** giraffe during the hot-dry (white), cold-dry (dark blue) and wet season (light blue) **B)** giraffe at different times of day: morning (light blue), midday (dark blue) and evening (white). Each boxplot for the three network metrics - i.e., scaled mean degree, scaled mean strength and transitivity - indicates the 95% confidence interval of the network metrics calculated for 100 random samples of the overall population. In addition, for each population, the real network metric estimates of the overall population networks are shown on the left of each of the plots - i.e., proportion equal to 100%. Animal silhouettes were downloaded from PhyloPic (https://www.phylopic.org/). Giraffe silhouette is by T. Michael Keesey under: Attribution-NonCommercial-ShareAlike 3.0 Unported (CC BY-NC-SA 3.0), link: https://creativecommons.org/licenses/by-nc-sa/3.0/.

**Figure 5:**
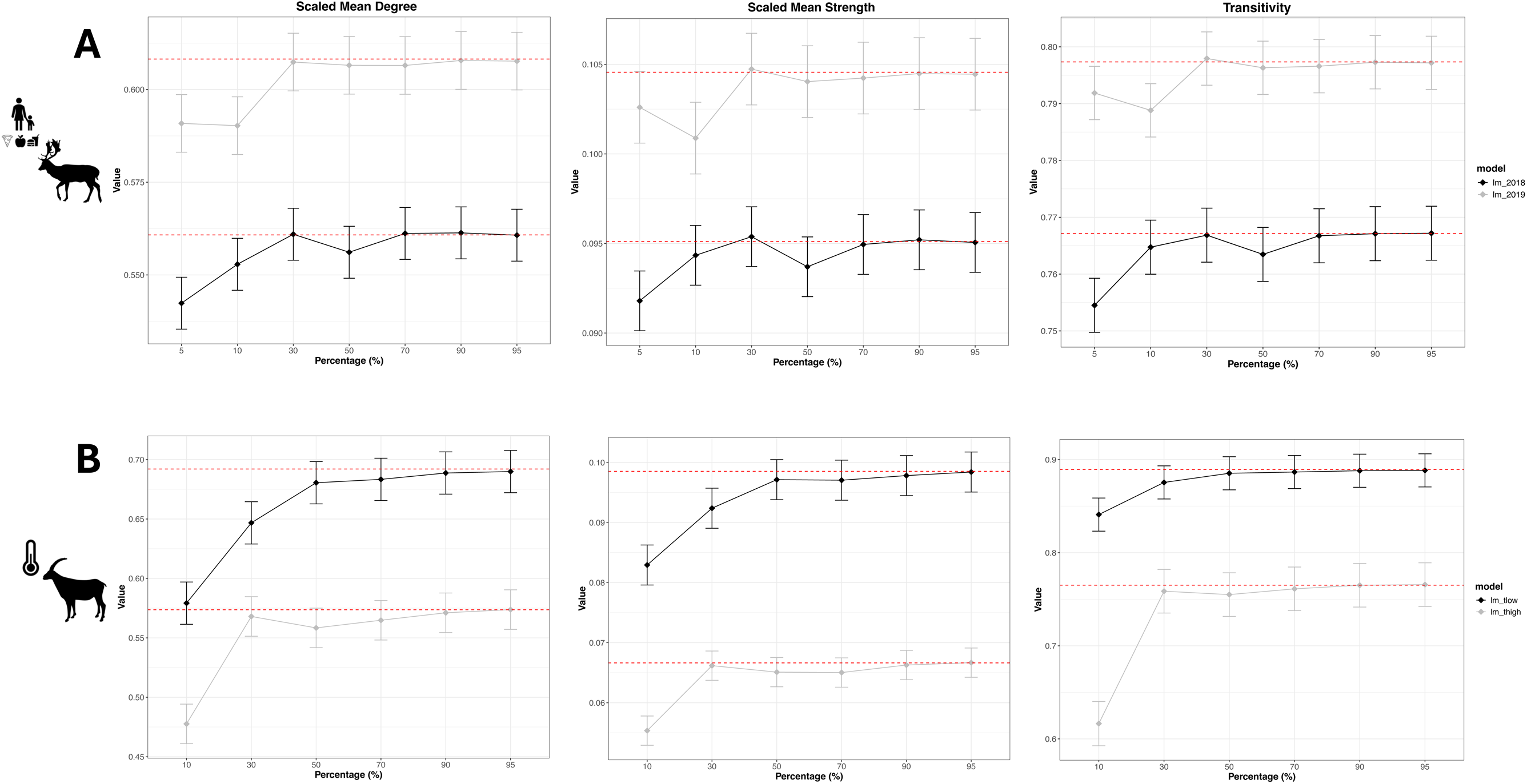
Plots showing the linear model estimates for each subsampling percentage and how they vary from the overall population networks (red dotted lines) for the three network metrics of interest - i.e., scaled mean degree, scaled mean strength and transitivity for the **A)** deer networks under human perturbation - i.e., grey indicating the subsamples of the 2018 social network and black of the 2019 social network; **B)** ibex networks under temperature extremes - i.e., grey indicating the subsamples of the low temperature social network and black of the high temperature social network. Animal silhouettes were downloaded from PhyloPic (https://www.phylopic.org/). Deer silhouette is by Anthony Caravaggi under: Attribution-NonCommercial-ShareAlike 3.0 Unported (CC BY-NC-SA 3.0), link: https://creativecommons.org/licenses/by-nc-sa/3.0/. Ibex silhouette is by Steven Traver under: CC0 1.0 Universal (CC0 1.0) Public Domain Dedication.

**Figure 6:**
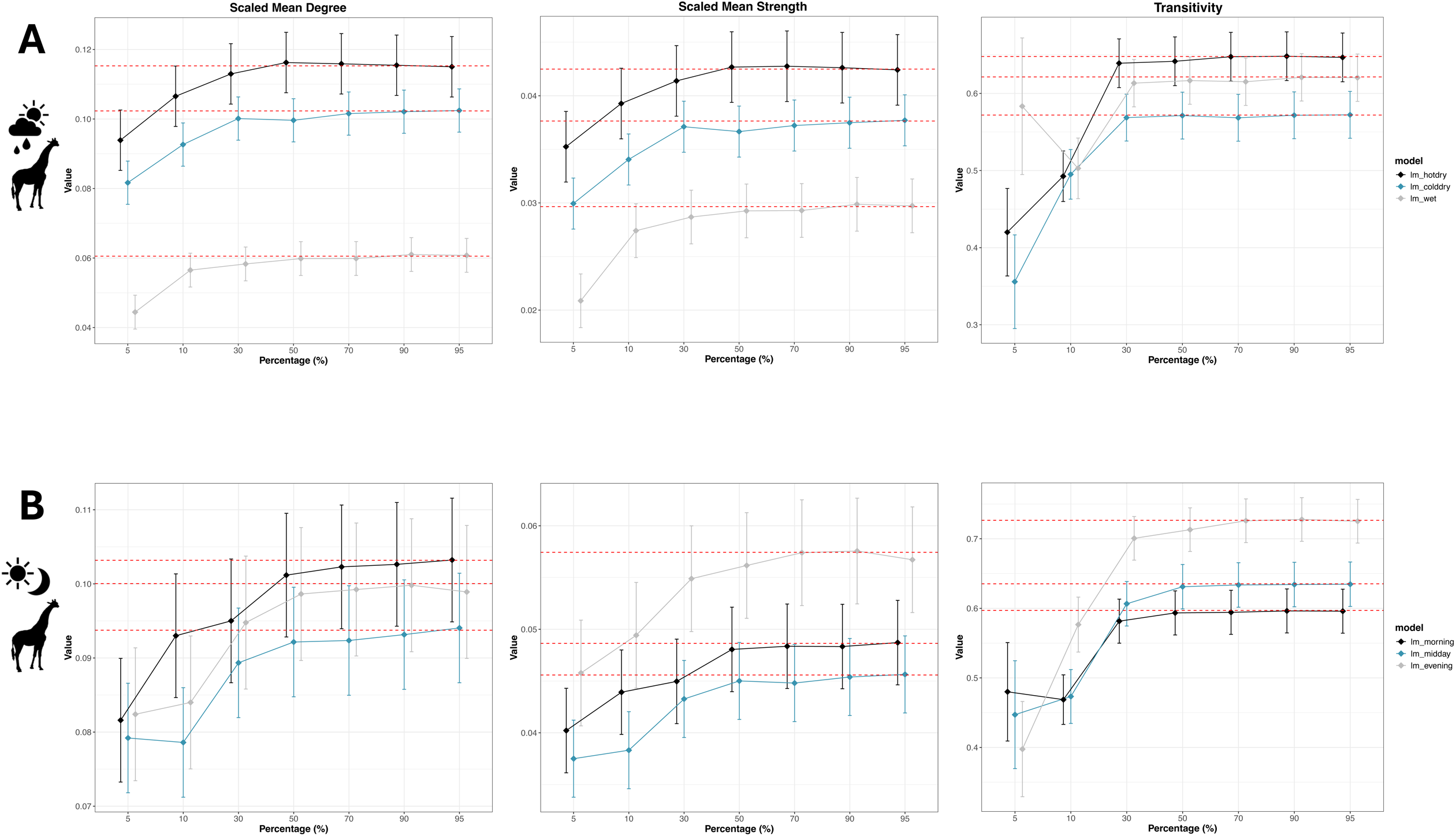
Plots showing the linear model estimates for each subsampling percentage and how they vary from the overall population networks (red dotted lines) for the three network metrics of interest - i.e., scaled mean degree, scaled mean strength and transitivity for the **A)** seasonal variation of the giraffe social networks - i.e., grey indicating the subsamples of the wet season social network, black of the hot-dry season social network and blue of the cold-dry season social network; **B)** daily variation (i.e., TOD - time of day) of the giraffe social networks - i.e., grey indicating the subsamples of the evening social network, black of the morning social network and blue of the midday social network. Animal silhouettes were downloaded from PhyloPic (https://www.phylopic.org/). Giraffe silhouette is by T. Michael Keesey under: Attribution-NonCommercial-ShareAlike 3.0 Unported (CC BY-NC-SA 3.0), link: https://creativecommons.org/licenses/by-nc-sa/3.0/.

**Figure 7:**
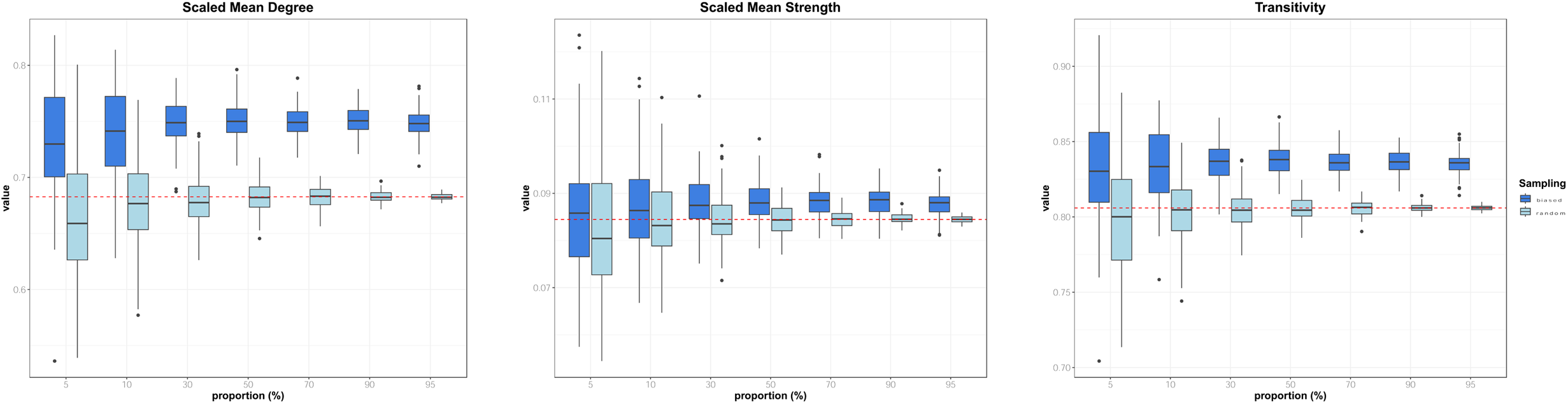
Plots showing the effect of random sampling (light blue; i.e., individuals randomly sampled from the overall population) compared to bias sampling (dark blue; i.e., increased probability of sampling “bold” individuals compared to “shy” individuals from the overall population). Each boxplot for the three network metrics - i.e., scaled mean degree, scaled mean strength and transitivity - indicates the 95% confidence interval of the network metrics calculated for 100 samples of the overall population. In addition, for each population, the real network metric estimates of the overall population network are indicated by a red dotted line.

### Goal 1: Effect of subsampling in comparing wildlife social network metrics

#### • Fallow deer: wildlife feeding

The 2019 deer networks (after reducing human-deer interactions, Figure 1B) had a higher degree and stronger connections compared to the networks recorded in 2018 (Figure 1A, before the management intervention), and this was the case for both main behavioural types present in the population (beggars or bold vs non-beggars or shy individuals). Social network metrics were higher in 2019 than in 2018 (Figure 3A), and such a statistically significant difference was maintained by random subsamples in the population including the 5% subsample (see Appendix S3 - Table S1 for the t-test results). However, only 30% of the population or higher random subsamples were able to provide a reliable estimate of all three network metrics that were not significantly different from the one recorded by the full population (Figure 5A - full details on linear models’ results presented in Appendix S4 - Table S5).

#### • Alpine ibex: temperature

The low temperature ibex network had a higher degree and stronger connections compared to the high temperature network (Figure 1C right *vs* left network). Social network metrics were higher at low temperatures compared to high temperatures (Figure 3B), and such a statistically significant difference was maintained by random subsamples in the population up to the 10% subsample (Appendix S3 - Table S2 for the t-test results). However, only 30% of the population or higher random subsamples were able to provide a reliable estimate of transitivity and 50% or higher for scaled mean degree and strength, which were not significantly different from the one recorded by the full population (Figure 5B - full details on linear models’ results presented in Appendix S4 - Table S6).

#### • Angolan giraffe: Season

Giraffe had a higher degree and stronger connections during the hot dry season compared to the other two seasons (Figure 2A), and such a statistically significant difference was maintained by random subsamples in the population up to the 10% subsample (Fig. 4A, See Appendix S3 - Table S3 for the t-test results). However, only 30% of the population or higher random subsamples were able to provide a reliable estimate of all three network metrics that were not significantly different from the one recorded by the full population (Figure 6A - full details on linear models’ results presented in Appendix S4 - Table S8).

#### • Angolan giraffe: TOD

Regarding diel changes in Angolan giraffe networks, the evening network had stronger connections compared to the other daily networks (Figure 2B). Scaled mean strength and transitivity were higher in the evening compared to the other TODs, whereas scaled mean degree was higher in the morning network (Figure 4B). These statistically significant differences were maintained by random subsamples in the population up to the 10% subsample for scaled mean strength and transitivity and 30% for scaled mean degree (See Appendix S3 - Table S4 for the t-test results). However, only 30% of the population or higher random subsamples were able to provide a reliable estimate of all three network metrics that were not significantly different from the one recorded by the full population (Figure 6B - full details on linear models’ results presented in Appendix S4 - Table S8).

### Goal 2: Effect of capture bias on social network metrics

In terms of random vs biased sampling by deer trappability, random sampling (nodes being sampled randomly, with their true probability; no bias towards a particular behavioural type: beggars P = 0.21, non-beggars P = 0.79) was compared to biased sampling (beggar nodes having a higher probability of being sampled, P = 0.91, compared to non-beggar nodes, P = 0.09) (Figure 7). When we used random sampling (light blue boxplot, Figure 7), population subsamples down to 10% reliably mimicked the full population social network metric estimate (red dotted line). Biased sampling, in contrast, generally resulted in over-estimating metrics (see detailed t-test results in Appendix S5 - Table S9).

## Discussion

In this study, we found intrinsic differences in the social networks of fallow deer, Alpine ibex and Angolan giraffe under human and climate pressures. Sociality of fallow deer was impacted by park visitors willing to feed them; high temperatures disintegrated ibex feeding aggregations; and a mixture of temperature and food availability, linked to seasonality and daily behavioural patterns shaped giraffe sociality in the Namib Desert. We demonstrated that random subsamples of such populations were still able to depict such differences in their sociality, showing that - as long as researchers have an accurate idea of population size - subsamples of wildlife populations can be carefully and efficiently used to make inferences on their sociality and provide empirical data for management, conservation and zoonotic disease ecology. Even small subsamples (e.g., 5%) of a natural population can depict intrinsic differences in sociality, however, such small subsamples cannot be used in most cases to estimate the actual network metrics of the entire population. In the case of fallow deer, for instance, a subsample of 5% correctly revealed the difference in sociality between the years before and after management intervention, but only subsamples equal to or greater than 30% could reveal the actual social network metrics estimated for the entire population. Therefore, depending on the research question (differences between networks vs estimating the actual network characteristics), researchers must aim accordingly for the appropriate sample size of monitored individuals. Despite the diverse species, ecological contexts and data collection regimes, we showed clear consistent patterns with minimal variation in subsample thresholds due to actual population size (e.g., smaller subsamples were more efficient in the larger population of deer compared to giraffe and ibex). Finally, our work underscores the importance of random sampling in wildlife populations. We showed that under-sampling of trap-shy individuals may introduce significant bias to animal network estimates. These results offer key guidelines to researchers using wildlife population subsamples when building social networks and studying changes in sociality, with clear implications for wildlife conservation and zoonotic disease transmission.

When using subsamples to compare social networks within the same population, we found differences in subsample performance depending on degrees of freedom (i.e., whether we compared two networks - such as in fallow deer and ibex - or three networks, in the case of giraffe). When comparing three networks, subsamples were less powerful in depicting differences in networks similar to the full population, suggesting larger sample sizes are needed when the number of levels increases. Similarly, smaller populations require researchers to boost the proportion of individuals monitored in a natural population. For example, 5% of the Ibex population (∼4 ibex out of a population of 65) is insufficient to provide any meaningful information compared to 5% of the fallow deer population (30 individuals out of a population of 600). Therefore, based on our results, 5% of the fallow deer population depicted differences between network metrics across years, although only 30% could provide unbiased full population network metric estimates. Whereas in giraffe (a large population but with three networks to be compared instead of two) and in ibex (a very small population), at least 10% of the individuals were necessary to depict differences across networks, whereas 30% (and in some cases 50% - e.g., mean strength in ibex) were needed to provide reliable estimates of the full network metrics.

When examining the effect of human disturbance on the behaviour of fallow deer, we found a significant difference in sociality before and after the management implementation aiming to reduce human-deer interactions [29]. When disturbed, herds in the park divide into smaller groups and are unable to settle into their normal social groups. This was due to some individuals (∼20%) staying to interact with humans (“bold”) while the others flee (“shy”), leading to a state of fluidity where the animals are unable to form stable groups and are constantly shifting and reorganizing [35]. These observations are consistent with the results shown by our social networks. After the management implementation in 2019 aimed at reducing human-deer interactions [29], we observed more and stronger connections between individuals, as well as higher clustering within the population, compared to 2018. This confirmed our prediction, demonstrating that animals can form stronger connections and more stable groups when disruptions are minimized. This further supports the success of the management implementation at lowering human disturbance levels, as already shown by Griffin *et al.* [29]. Understanding the effect of human disturbances and management solutions on animal behaviour is essential to inform conservation efforts and policies [15, 40]. Alterations to natural behaviour due to humans (e.g., directly interacting with wildlife, altering natural habitats for agriculture or residential development) can result in adaptations of feeding patterns, reduced reproductive success and increased stress [15, 40], which have been shown to have direct effects on animal fitness and survival [41–43]. Therefore, monitoring changes in animal behaviour with reliable analytical tools allows us to implement effective measures that promote coexistence between humans and wildlife, ensuring their mutual welfare.

In addition, we were interested in exploring how animal sociality is influenced by weather and climate factors. Our findings indicated that ibex were more clustered and had more and stronger connections when the temperatures were < 10 °C compared to > 14 °C. Ibex are highly susceptible to temperature extremes, needing to feed in higher pastures, but also suboptimal in quality and quantity, to avoid overheating during days with high temperatures, while they tend to form bigger groups and graze all together when the temperatures drop and they can use pastures with higher quality/quantity of forage [30]. Similarly, giraffe were found to be more clustered and have more and stronger connections during the dry seasons relative to the wet season, consistent with our *a priori* hypothesis. During the dry season, food is less readily available, forcing giraffe to travel long distances and cluster in areas with higher food availability [44, 45]. Moreover, we observed a relationship between time of day and giraffe sociality: giraffe were more clustered and had stronger connections in the evening when they sought shelter and prepared for nocturnal predation threats [31, 46]. In contrast, individuals exhibited more social interactions in the morning while foraging and interacting with other groups. This behaviour was consistent with previous studies [31, 47, 48], as well as our predictions. Shifts in spatiotemporal animal behaviour patterns resulting from anthropogenic climate change can be high-impact, leading to reduced resource availability, lower reproductive success and metabolic disruptions (e.g., altered energy budgets due to greater thermoregulatory investment) [49–51]. Hence, understanding the full ethological ramifications of changes in key climatic factors can enhance the efficacy of related wildlife conservation strategies, and thus we recommend future research be conducted in this direction.

Zoonotic disease transmission is a fundamental factor to consider when investigating changes in animal behaviour linked to human perturbation and climate change pressures. A clear example is the observed sociality shift in the peri-urban fallow deer population following the implementation of specific management actions which led to a significant reduction in human-deer interactions [29], resulting in an increase in transitivity within the fallow deer social network. Transitivity is negatively correlated with disease transmission [52], as higher transitivity indicates greater clustering of individuals. This clustering means that if one cluster is infected, other clusters remain relatively isolated and protected due to limited connections between them. However, continuous human disruptions can increase disease transmission by enhancing connectivity between clusters as groups constantly shift and reorganize. Therefore, understanding how human perturbations alter animal social dynamics is crucial for predicting and managing zoonotic disease transmission. This is particularly important given that COVID-19 has been found in deer populations, likely spreading through interactions with humans [53]. In addition, changes in climate, such as rising temperatures and altered seasonal patterns, can cause animals to aggregate due to the concentration of resources in fewer locations and the decreased availability of suitable shelters for thermoregulation [54]. Aggregation of individuals reduces network transitivity by increasing interactions between groups, which leads to higher disease transmission. For example, our findings indicate the ibex population exhibited less clustering at higher temperatures, increasing the probability of disease transmission. Given the current climate crisis and the rise in human-wildlife interactions, understanding how these factors influence animal sociality is pivotal in accurately forecasting the transmission of zoonoses and predicting threats to emergently vulnerable species.

Our work illustrated how and when random subsamples can be used to make population-wide inferences and reveal intrinsic differences within population networks. However, we also demonstrated that non-random sampling (e.g., sampling affected by trappability of individuals) can significantly bias social network estimates. In the fallow deer population, for instance, the vast majority of newborns are tagged during the first week(s) of life [55], but only ∼20% of them display the tendency to approach humans and accept food from their hands. Hence, these would be the first animals that any researchers would be able to immobilise and tag for longitudinal research. Therefore, when using social networks to study behavioural changes, the effects of behaviour variability within the population must be considered [25, 26]. Animals have been widely shown to display consistent and repeatable interpersonal variability in the way they behave - i.e., animal personality [56]. Nonetheless, animal personality has been underexploited in ecological research, particularly when making inferences about entire populations from subsamples. Indeed, previous research has shown a capture bias towards individuals that show more risky behaviours (“bold”) [25], a bias whose effects on social network metrics relative to random sampling were previously unexplored. Herein we observed significant resultant bias in the fallow deer population network when we increased the probability to sample individuals that consistently beg for food. Networks that were biased towards bolder individuals were shown to be more clustered as well as having more and stronger connections. This pattern held true for all subsamples except for scaled mean strength, which did not significantly differ from random sampling when subsamples were very small (5%). Understanding how this bias influences networks is crucial, as it can significantly skew our inferences. Therefore, obtaining robust estimates by randomly sampling animals is essential in calculating reliable social networks that can be used for effective conservation management.

In conclusion, social network analysis is a powerful tool for understanding how animals adapt to changing environments. Collecting comprehensive behavioural and movement data on entire populations to construct social networks is often impractical [16]. Our study demonstrates that population subsamples can reveal intrinsic differences in animal behaviour effectively, provided they are both randomly sampled and sufficiently large to produce reliable estimates, without which targeted wildlife conservation and prediction of zoonosis transmissions cannot be achieved. Future research should therefore focus on leveraging global databases and conducting simulations to determine the percentage of the population needed for reliable estimates and how it is affected by the total population sample size. Moreover, new research should aim to improve our understanding and prediction of how climate change and human disturbances impact animal sociality and zoonotic disease transmission, as well as identify effective management strategies.

## Acknowledgements

This publication has emanated from research conducted with the financial support of Science Foundation Ireland under Grant number 18/CRT/6049. For the purpose of Open Access, the author has applied a CC BY public copyright licence to any Author Accepted Manuscript version arising from this submission. SG was supported by FAR 2023-2024 and FIRD 2023 of the University of Ferrara.

## Author Contributions

**KC:** conceptualization, data curation, formal analysis, writing—original draft, writing—review and editing.

**PK:** formal analysis, writing—review and editing.

**MB, JF, SF, EH:** data curation (Angolan giraffe), writing—review and editing.

**BA, AD, LLG, JF, AH, LK:** data curation (fallow deer), writing—review and editing.

**SG, FB:** data curation (Alpine ibex), writing—review and editing.

**MST:** formal analysis, supervision, writing—review and editing.

**SC:** conceptualization, formal analysis, supervision, writing—review and editing.

All authors gave final approval for publication and agreed to be held accountable for the work performed therein.

