## Appendices for "Random subsamples of animal populations can reveal intrinsic differences in sociality with key implications in ecology, conservation and disease transmission"

**Appendix S1**

- *Fallow deer population*

The first population that we included in this study is a fallow deer (*Dama dama*) population residing in the largest enclosed urban park in Europe; Phoenix Park, located in Dublin, IE (53°22'N, -6°21'W). The park is 709 ha and has three main habitats: 56% grassland, 31% woodland; and 7% road and buildings [1]. The population is maintained at ~600 individuals (end of year population estimates, OPW official data) by the park management with yearly culls. Females and males are sexually segregated for most of the year (except during the rutting season between end of September and the beginning of November). Newborn male fawns spend the first 1-2 years of their lives in the female sector, after which they leave to join bachelor groups in the male sector.

The dataset used in this study comprises of focal observations conducted between May 2018 and January 2023. The data collection is conducted all year round with weekly frequency between August and May, and a more frequent collection during the remaining months (5 days a week between 2018-20, 2 days a week between 2021-23) due to a seasonal study on human-wildlife conflict run during these months [2]. For the collection the park is divided into sectors with a stratified *a priori* schedule which ensures the park sectors are equally sampled. Sectors are walked systematically and all groups of fallow deer are recorded (i.e., a group is defined as 2 or more deer being less than 50m apart and within view of each other). For each group observation the unique animal IDs, date, time, and location is collated and given a unique group ID [3]. Individuals are identified by unique ID ear tags which are applied in their first 1-3 weeks of life (~85% of the population is tagged) [4]. Quality control on observational data to remove rare “ghost” animals (i.e., animals with unofficial names, likely misidentifications) was conducted by using only those animals whose IDs were officially recorded or were observed across ≥2 sets of ≥2 consecutive sampling months 2018-23. The total number of animals used in our final network looking at the effect of management between 2018 and 2019 was 550 individuals for both years (~85% of the population). As for our analysis on the entire population between May 2018 and January 2023 we used 751 individuals (which were classified in behavioural categories and estimated to be ~70% of the population, see [5])

- *Alpine ibex population*

The second population included in the study is an ibex (*Capra ibex*) population that resides in the Gran Paradiso National Park in the north-west of Italy (45°35′N, 7°12′E). The population lives in the Levionaz valley which is around 1700 ha, with an altitude between 1700 and 2300m a.s.l. (above sea level). Below 2300m a.s.l., the habitat is mainly characterised by meadows and conifer patches, while at higher altitudes it is composed mainly of rocks, meadows and grassland [6].

Observational data used from this population was collected between 2010-2011 (early May to late October) on male ibex only (i.e., male population estimated to be 51 and 60 individuals with 44 and 52 marked individuals for 2010 and 2011, respectively) [6]. Observations therefore were obtained for ~86% percent of the male ibex population (57 total tagged individuals between the two years, with an estimated total population size of 66 unique individuals) living in the study area [6], which are identifiable by their unique ear tags (captured when subadult and adult, ≥ 2 years old, by means of chemical immobilization, see [7]). Each observation included a unique group ID, date, time, maximum daily temperature as well as the individual animal IDs observed. The total number of animals used in our final network was of 51 individuals (~80% of the population).

- *Angolan giraffe population*

The third population used in the study is an Angolan giraffe (*Giraffa giraffa angolensis*) population that resides in the northern Namib Desert in northwest Namibia (− 18.94661°N, 13.05204°E). The region is characterised by an ephemeral river system as well as a seasonal climate which consists of three seasons: cold-dry (June-August), hot-dry (September-February), wet-dry (March-May) [8].

The data collection for this population was run between February 2016 and July 2019 [8, 9]. It was conducted from a vehicle, with a total of 34 sampling periods, ranging between 7 to 15 days. This variation was due to unforeseeable circumstances (e.g., vehicle breakdown etc.). The collection was run around a 250km transect. Sampling was carried out in all sectors for each sampling period, with the starting location and direction of travel changed systematically to reduce sampling bias. Each group of giraffe observed had a unique group ID, location, date, time and its composition (i.e., age, sex, and individual IDs) [8, 9]. Individuals are identified by their unique pelage patterns, which are also verified using *HotSpotter* [10] (~95% of the total population was observed during the sampling period). In each sampling day a group was recorded only if none of the individuals in the group had already been observed on the same day to avoid pseudo-replication. In the final dataset of this species we also excluded all individuals with 5 or less observations (i.e., during the total collection period of 3.5 years), < 1 year old (i.e., juveniles have similar social encounters from their mothers), only sighted in the far north of the study site (i.e., less evenly sampled due to logistical reasons) [11]. The total population estimate living in the south study site is ~350 individuals (i.e., of which 334 individuals were observed). The total number of animals used in our final network was of 123 individuals for the season networks (~35% of the total population) and 104 for the time of day networks (~30% of the total population).

**Appendix S2**

Here we compare 200 samples of the population where in 100 of the subsamples 158 nodes, consisting of bold individuals (consistent beggars) of the population, are picked with replacement (purple) and in the other 100 samples they are selected without replacement (yellow). As for the shy individuals (occasional beggars and avoiders) 158 nodes are always picked randomly for all the 200 samples, both with (purple) and without (yellow) replacement. We used 158 nodes for each behavioral type since it was the highest amount of nodes we could use to be able to select bold individuals without replacement (i.e., the population has 158 individuals that consistently beg for food from human, a.k.a bold individuals). We performed this analysis to ensure that sampling with and without replacement would not affect the analysis where we compare the effect of capture bias on network metrics. All three metrics show to not be significantly different between samples with and without replacement.

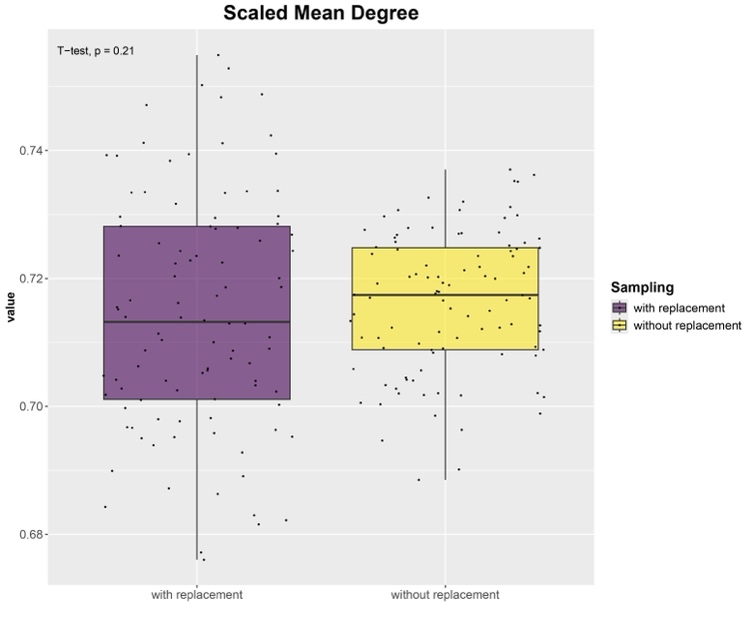

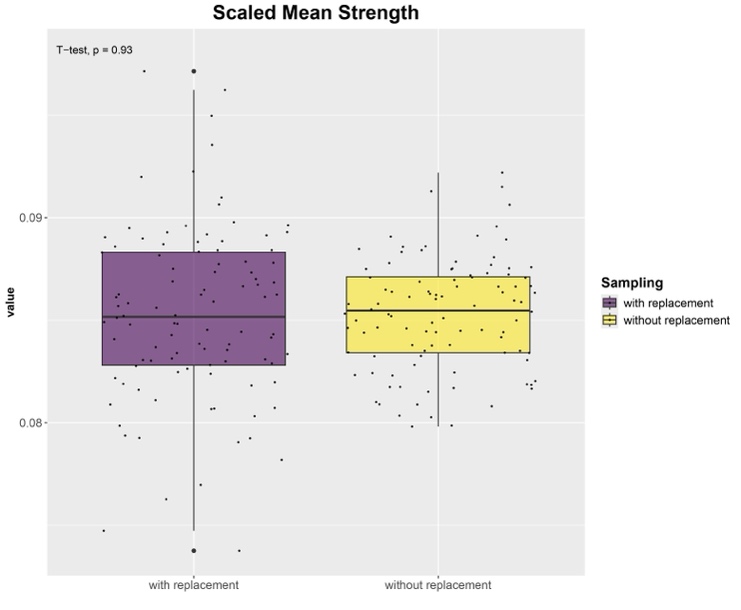

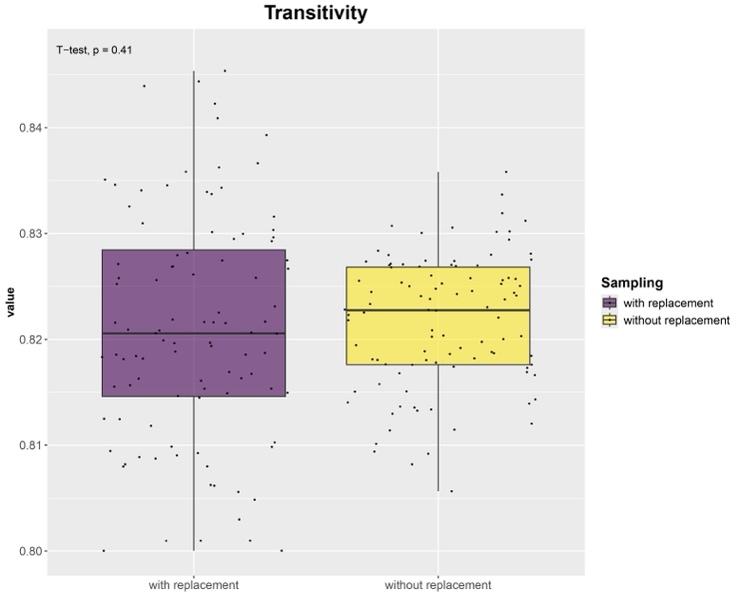

**Figure S1:** Comparison of sampling without replacement (yellow) and with replacement (purple). The paired t-test p-value is shown on all plots on the top-left corner.

**Appendix S3**

**Table S1: Tables of the unpaired t-tests for the fallow deer population – Year 2018 VS 2019**

| Subsample % | Welch Two sample t-test | Metric | | |
| --- | --- | --- | --- | --- |
|  |  | Scaled Mean Degree | Scaled Mean Strength | Transitivity |
| 95% | 2018 | 0.561 | 0.095 | 0.767 |
|  | 2019 | 0.608 | 0.104 | 0.797 |
|  | P-value | **< 0.001** *** | **< 0.001** *** | **< 0.001** *** |
|  | T-statistics | -81.306 | -67.220 | -89.746 |
| 90% | 2018 | 0.561 | 0.095 | 0.767 |
|  | 2019 | 0.608 | 0.104 | 0.797 |
|  | P-value | **< 0.001** *** | **< 0.001** *** | **< 0.001** *** |
|  | T-statistics | -52.321 | -46.396 | -60.485 |
| 70% | 2018 | 0.561 | 0.095 | 0.767 |
|  | 2019 | 0.606 | 0.104 | 0.797 |
|  | P-value | **< 0.001** *** | **< 0.001** *** | **< 0.001** *** |
|  | T-statistics | -29.674 | -24.494 | -31.851 |
| 50% | 2018 | 0.556 | 0.094 | 0.763 |
|  | 2019 | 0.607 | 0.104 | 0.796 |
|  | P-value | **< 0.001** *** | **< 0.001** *** | **< 0.001** *** |
|  | T-statistics | -18.657 | -15.093 | -19.704 |
| 30% | 2018 | 0.561 | 0.095 | 0.767 |
|  | 2019 | 0.607 | 0.105 | 0.798 |
|  | P-value | **< 0.001** *** | **< 0.001** *** | **< 0.001** *** |
|  | T-statistics | -11.450 | -9.120 | -12.808 |
| 10% | 2018 | 0.553 | 0.094 | 0.765 |
|  | 2019 | 0.590 | 0.101 | 0.789 |
|  | P-value | **< 0.001** *** | **< 0.001** *** | **< 0.001** *** |
|  | T-statistics | -4.798 | -3.480 | -4.950 |
| 5% | 2018 | 0.542 | 0.092 | 0.755 |
|  | 2019 | 0.591 | 0.103 | 0.792 |
|  | P-value | **< 0.001** *** | **< 0.001** *** | **< 0.001** *** |
|  | T-statistics | -4.602 | -4.066 | -5.424 |
|  | Note: P-value: 0.1-1 **NS**, 0.05 *****, 0.01 ******, 0.001 ******* | | | |

**Table S2: Tables of the unpaired t tests for the Alpine ibex population - Temperature low VS high**

| Subsample % | Welch Two sample t-test | Metric | | |
| --- | --- | --- | --- | --- |
|  |  | Scaled Mean Degree | Scaled Mean Strength | Transitivity |
| 95% | < 10 °C | 0.690 | 0.098 | 0.888 |
|  | > 14 °C | 0.575 | 0.067 | 0.766 |
|  | P-value | **< 0.001** *** | **< 0.001** *** | **< 0.001** *** |
|  | T-statistics | 51.204 | 86.549 | 102.614 |
| 90% | < 10 °C | 0.689 | 0.098 | 0.888 |
|  | > 14 °C | 0.571 | 0.067 | 0.763 |
|  | P-value | **< 0.001** *** | **< 0.001** *** | **< 0.001** *** |
|  | T-statistics | 32.234 | 51.720 | 61.603 |
| 70% | < 10 °C | 0.683 | 0.098 | 0.887 |
|  | > 14 °C | 0.566 | 0.066 | 0.759 |
|  | P-value | **< 0.001** *** | **< 0.001** *** | **< 0.001** *** |
|  | T-statistics | 17.862 | 29.222 | 36.567 |
| 50% | < 10 °C | 0.681 | 0.098 | 0.885 |
|  | > 14 °C | 0.553 | 0.065 | 0.749 |
|  | P-value | **< 0.001** *** | **< 0.001** *** | **< 0.001** *** |
|  | T-statistics | 12.192 | 18.971 | 22.105 |
| 30% | < 10 °C | 0.647 | 0.092 | 0.875 |
|  | > 14 °C | 0.559 | 0.066 | 0.749 |
|  | P-value | **< 0.001** *** | **< 0.001** *** | **< 0.001** *** |
|  | T-statistics | 6.159 | 11.107 | 13.958 |
| 10% | < 10 °C | 0.579 | 0.083 | 0.841 |
|  | > 14 °C | 0.505 | 0.057 | 0.666 |
|  | P-value | **0.002** ** | **< 0.001** *** | **< 0.001** *** |
|  | T-statistics | 3.138 | 6.134 | 5.132 |
|  | Note: P-value: 0.1-1 **NS**, 0.05 *****, 0.01 ******, 0.001 ******* | | | |

**Table S3: Tables of the linear models for the Angolan giraffe population - Seasons**

| Subsample % | Season | Scaled Mean Degree | | Scaled Mean Strength | | Transitivity | |
| --- | --- | --- | --- | --- | --- | --- | --- |
|  |  | Estimate | Std. Error | Estimate | Std. Error | Estimate | Std. Error |
| 95% | (Intercept) | 0.102 *** | 0.0003432 | 0.038 *** | 0.0001229 | 0.572 *** | 0.0009407 |
|  | Hot-Dry | 0.013 *** | 0.0004853 | 0.005 *** | 0.0001738 | 0.074 *** | 0.0013304 |
|  | Wet | -0.042 *** | 0.0004853 | -0.008 *** | 0.0001738 | 0.048 *** | 0.0013304 |
| 90% | (Intercept) | 0.102 *** | 0.0004872 | 0.037 *** | 0.0001774 | 0.572 *** | 0.001329 |
|  | Hot-Dry | 0.013 *** | 0.0006890 | 0.005 *** | 0.0002508 | 0.077 *** | 0.001879 |
|  | Wet | -0.041 *** | 0.0006890 | -0.008 *** | 0.0002508 | 0.049 *** | 0.001879 |
| 70% | (Intercept) | 0.102 *** | 0.0008899 | 0.037 *** | 0.0003142 | 0.568 *** | 0.002786 |
|  | Hot-Dry | 0.014 *** | 0.0012586 | 0.006 *** | 0.0004444 | 0.079 *** | 0.003940 |
|  | Wet | -0.042 *** | 0.0012586 | -0.008 *** | 0.0004444 | 0.047 *** | 0.003940 |
| 50% | (Intercept) | 0.100 *** | 0.001332 | 0.037 *** | 0.0004866 | 0.571 *** | 0.004248 |
|  | Hot-Dry | 0.017 *** | 0.001883 | 0.006 *** | 0.0006882 | 0.070 *** | 0.006008 |
|  | Wet | -0.040 *** | 0.001883 | -0.007 *** | 0.0006882 | 0.045 *** | 0.006008 |
| 30% | (Intercept) | 0.100 *** | 0.002270 | 0.037 *** | 0.0008306 | 0.569 *** | 0.009284 |
|  | Hot-Dry | 0.013 *** | 0.003211 | 0.004 *** | 0.0011746 | 0.071 *** | 0.013130 |
|  | Wet | -0.042 *** | 0.003211 | -0.008 *** | 0.0011746 | 0.045 *** | 0.013130 |
| 10% | (Intercept) | 0.093 *** | 0.004832 | 0.034 *** | 0.001954 | 0.495 *** | 0.036311 |
|  | Hot-Dry | 0.014 ***** | 0.006833 | 0.005 **NS** | 0.002764 | -0.002 **NS** | 0.050796 |
|  | Wet | -0.036 *** | 0.006833 | -0.007 * | 0.002764 | 0.008 **NS** | 0.056940 |
| 5% | (Intercept) | 0.082 *** | 0.00721 | 0.030 *** | 0.002985 | 0.356 *** | 0.08845 |
|  | Hot-Dry | 0.012 **NS** | 0.01020 | 0.005 **NS** | 0.004221 | 0.064 **NS** | 0.11889 |
|  | Wet | -0.037 *** | 0.01020 | -0.009 ***** | 0.004221 | 0.228 **NS** | 0.15532 |
|  | Note: P-value: 0.1-1 **NS**, 0.05 *****, 0.01 ******, 0.001 *******  Estimates: Indicate the difference in estimates at each subsample size compared to the Cold-Dry season estimate | | | | | | |

**Table S4: Tables of the linear models for the Angolan giraffe population - TOD**

| Subsample % | TOD | Scaled Mean Degree | | Scaled Mean Strength | | Transitivity | |
| --- | --- | --- | --- | --- | --- | --- | --- |
|  |  | Estimate | Std. Error | Estimate | Std. Error | Estimate | Std. Error |
| 95% | (Intercept) | 0.100 *** | 0.0003981 | 0.058 *** | 0.0001997 | 0.727 *** | 0.001023 |
|  | Midday | -0.007 *** | 0.0005631 | -0.012 *** | 0.0002824 | -0.090 *** | 0.001447 |
|  | Morning | 0.003 *** | 0.0005631 | -0.009 *** | 0.0002824 | -0.130 *** | 0.001447 |
| 90% | (Intercept) | 0.099 *** | 0.0005474 | 0.057 *** | 0.0002738 | 0.725 *** | 0.001472 |
|  | Midday | -0.005 *** | 0.0007741 | -0.011 *** | 0.0003872 | -0.091 *** | 0.002082 |
|  | Morning | 0.003 *** | 0.0007741 | -0.009 *** | 0.0003872 | -0.127 *** | 0.002082 |
| 70% | (Intercept) | 0.099 *** | 0.001136 | 0.057 *** | 0.0005585 | 0.724 *** | 0.002935 |
|  | Midday | -0.006 *** | 0.001606 | -0.012 *** | 0.0007898 | -0.087 *** | 0.004151 |
|  | Morning | 0.002 **NS** | 0.001606 | -0.009 *** | 0.0007898 | -0.130 *** | 0.004151 |
| 50% | (Intercept) | 0.098 *** | 0.001646 | 0.056 *** | 0.000771 | 0.719 *** | 0.004521 |
|  | Midday | -0.004 **NS** | 0.002328 | -0.011 *** | 0.001090 | -0.089 *** | 0.006394 |
|  | Morning | 0.004 **NS** | 0.002328 | -0.008 *** | 0.001090 | -0.123 *** | 0.006394 |
| 30% | (Intercept) | 0.091 *** | 0.002442 | 0.052 *** | 0.001224 | 0.694 *** | 0.008686 |
|  | Midday | 0.001 **NS** | 0.003453 | -0.007 *** | 0.001731 | -0.072 *** | 0.012284 |
|  | Morning | 0.009 ***** | 0.003453 | -0.005 ** | 0.001731 | -0.111 *** | 0.012284 |
| 10% | (Intercept) | 0.096 *** | 0.00606 | 0.053 *** | 0.002995 | 0.585 *** | 0.04493 |
|  | Midday | -0.012 **NS** | 0.00857 | -0.012 ** | 0.004236 | -0.117 **NS** | 0.06267 |
|  | Morning | -0.002 **NS** | 0.00857 | -0.010 * | 0.004236 | -0.162 ** | 0.06208 |
| 5% | (Intercept) | 0.098 *** | 0.009753 | 0.053 *** | 0.004769 | 0.350 *** | 0.08708 |
|  | Midday | -0.010 **NS** | 0.013793 | -0.013 **NS** | 0.006745 | -0.028 **NS** | 0.13697 |
|  | Morning | -0.006 **NS** | 0.013793 | -0.013 **NS** | 0.006745 | 0.122 **NS** | 0.12818 |
|  | Note: P-value: 0.1-1 **NS**, 0.05 *****, 0.01 ******, 0.001 *******  Estimates: Indicate the difference in estimates at each subsample size compared to the Evening TOD estimate | | | | | | |

**Appendix S4**

**Tables showing the summary for the linear models looking at the effect of subsampling on the network metric estimates for all three populations.**

**Table S5: Fallow deer – Year 2018 VS 2019**

| Level | Subsample % | Scaled Mean Degree | | Scaled Mean Strength | | Transitivity | |
| --- | --- | --- | --- | --- | --- | --- | --- |
|  |  | Estimate | Std. Error | Estimate | Std. Error | Estimate | Std. Error |
| 2018 | (Intercept) | 0.561 ******* | 0.004 | 0.095 ******* | 0.0008 | 0.767 ******* | 0.002 |
|  | 90% | 0.0006 **NS** | 0.005 | 0.0001 **NS** | 0.0012 | -8.113^-5^ **NS** | 0.003 |
|  | 70% | 0.0005 **NS** | 0.005 | -0.0001 **NS** | 0.0012 | -0.0004 **NS** | 0.003 |
|  | 50% | -0.005 **NS** | 0.005 | -0.0014 **NS** | 0.0012 | -0.0037 **NS** | 0.003 |
|  | 30% | 0.0003 **NS** | 0.005 | 0.0003 **NS** | 0.0012 | -0.0003 **NS** | 0.003 |
|  | 10% | -0.008 **NS** | 0.005 | -0.0007 **NS** | 0.0012 | -0.0025 **NS** | 0.003 |
|  | 5% | -0.018 ******* | 0.005 | -0.0033 ****** | 0.0012 | -0.013 ******* | 0.003 |
| 2019 | (Intercept) | 0.608 ******* | 0.004 | 0.105 ******* | 0.0010 | 0.797 ******* | 0.002 |
|  | 90% | 0.0002 **NS** | 0.006 | 3.641^-5^ **NS** | 0.0014 | 0.0001 **NS** | 0.003 |
|  | 70% | -0.0012 **NS** | 0.006 | -2.143^-4^ **NS** | 0.0014 | -0.0006 **NS** | 0.003 |
|  | 50% | -0.0011 **NS** | 0.006 | -4.109^-4^ **NS** | 0.0014 | -0.0009 **NS** | 0.003 |
|  | 30% | -0.0002 **NS** | 0.006 | 2.865^-4^ **NS** | 0.0014 | 0.0008 **NS** | 0.003 |
|  | 10% | -0.0174 ** | 0.006 | -3.566^-3^ * | 0.0014 | -0.0084 ***** | 0.003 |
|  | 5% | -0.0168 ** | 0.006 | -1.846^-3^ **NS** | 0.0014 | -0.0053 **NS** | 0.003 |
|  | Note: P-value: 0.1-1 **NS**, 0.05 *****, 0.01 ******, 0.001 *******  Estimates: Indicate the difference in estimates at each subsample size compared to the 95%  estimate | | | | | | |

**Table S6: Alpine Ibex – Temperature low VS high**

| Level | Subsample % | Scaled Mean Degree | | Scaled Mean Strength | | Transitivity | |
| --- | --- | --- | --- | --- | --- | --- | --- |
|  |  | Estimate | Std. Error | Estimate | Std. Error | Estimate | Std. Error |
| < 10 °C | (Intercept) | 0.690 ******* | 0.009 | 0.098 ******* | 0.002 | 0.888 ******* | 0.009 |
|  | 90% | -0.001 **NS** | 0.013 | -0.0006 **NS** | 0.002 | -0.0004 **NS** | 0.013 |
|  | 70% | -0.007 **NS** | 0.013 | -0.0014 **NS** | 0.002 | -0.0018 **NS** | 0.013 |
|  | 50% | -0.009 **NS** | 0.013 | -0.0013 **NS** | 0.002 | -0.003 **NS** | 0.013 |
|  | 30% | -0.043 ******* | 0.013 | -0.006 ***** | 0.002 | -0.013 **NS** | 0.013 |
|  | 10% | -0.111 ******* | 0.013 | -0.015 ******* | 0.002 | -0.047 ******* | 0.013 |
| > 14 °C | (Intercept) | 0.574 ******* | 0.008 | 0.067 ******* | 0.001 | 0.766 ******* | 0.012 |
|  | 90% | -0.003 **NS** | 0.012 | -0.0004 **NS** | 0.002 | -0.0007 **NS** | 0.017 |
|  | 70% | -0.009 **NS** | 0.012 | -0.002 **NS** | 0.002 | -0.005 **NS** | 0.017 |
|  | 50% | -0.015 **NS** | 0.012 | -0.002 **NS** | 0.002 | -0.011 **NS** | 0.017 |
|  | 30% | -0.006 **NS** | 0.012 | -0.0005 **NS** | 0.002 | -0.007 **NS** | 0.017 |
|  | 10% | -0.096 ******* | 0.012 | -0.011 ******* | 0.002 | -0.149 ******* | 0.017 |
|  | Note: P-value: 0.1-1 **NS**, 0.05 *****, 0.01 ******, 0.001 *******  Estimates: Indicate the difference in estimates at each subsample size compared to the 95%  estimate | | | | | | |

**Table S7: Angolan giraffe - Season**

| Level | Subsample % | Scaled Mean Degree | | Scaled Mean Strength | | Transitivity | |
| --- | --- | --- | --- | --- | --- | --- | --- |
|  |  | Estimate | Std. Error | Estimate | Std. Error | Estimate | Std. Error |
| Hot-Dry | (Intercept) | 0.115 ******* | 0.004 | 0.042 ******* | 0.002 | 0.647 *** | 0.016 |
|  | 90% | 0.0004 **NS** | 0.006 | 0.0002 **NS** | 0.002 | 0.0015 **NS** | 0.023 |
|  | 70% | 0.0009 **NS** | 0.006 | 0.0003 **NS** | 0.002 | 0.0009 **NS** | 0.023 |
|  | 50% | 0.001 **NS** | 0.006 | 0.0003 **NS** | 0.002 | -0.005 **NS** | 0.023 |
|  | 30% | -0.002 **NS** | 0.006 | -0.0010 **NS** | 0.002 | -0.008 **NS** | 0.023 |
|  | 10% | -0.008 **NS** | 0.006 | -0.0031 **NS** | 0.002 | -0.154 ******* | 0.023 |
|  | 5% | -0.021 ******* | 0.006 | -0.0072 ****** | 0.002 | -0.227 ******* | 0.033 |
| Cold-Dry | (Intercept) | 0.102 ******* | 0.003 | 0.038 ******* | 0.001 | 0.572 ******* | 0.015 |
|  | 90% | -0.0003 **NS** | 0.004 | -0.0002 **NS** | 0.002 | -0.0006 **NS** | 0.022 |
|  | 70% | -0.0009 **NS** | 0.004 | -0.0005 **NS** | 0.002 | -0.0038 **NS** | 0.022 |
|  | 50% | -0.0028 **NS** | 0.004 | -0.0011 **NS** | 0.002 | -0.001 **NS** | 0.022 |
|  | 30% | -0.0023 **NS** | 0.004 | -0.0006 **NS** | 0.002 | -0.004 **NS** | 0.022 |
|  | 10% | -0.0098 ***** | 0.004 | -0.0037 ***** | 0.002 | -0.077 ******* | 0.023 |
|  | 5% | -0.002 ******* | 0.004 | -0.008 ******* | 0.002 | -0.216 ******* | 0.035 |
| Wet | (Intercept) | 0.061 ******* | 0.002 | 0.030 ******* | 0.001 | 0.620 ******* | 0.016 |
|  | 90% | 0.0002 **NS** | 0.004 | 0.0001 **NS** | 0.002 | 0.0005 **NS** | 0.022 |
|  | 70% | -0.0009 **NS** | 0.004 | -0.0004 **NS** | 0.002 | -0.005 **NS** | 0.022 |
|  | 50% | -0.0009 **NS** | 0.004 | -0.0005 **NS** | 0.002 | -0.004 **NS** | 0.022 |
|  | 30% | -0.0025 **NS** | 0.004 | -0.001 **NS** | 0.002 | -0.007 **NS** | 0.022 |
|  | 10% | -0.0042 **NS** | 0.004 | -0.002 **NS** | 0.002 | -0.118 ******* | 0.025 |
|  | 5% | -0.016 ******* | 0.004 | -0.009 ******* | 0.002 | -0.037 **NS** | 0.048 |
|  | Note: P-value: 0.1-1 **NS**, 0.05 *****, 0.01 ******, 0.001 *******  Estimates: Indicate the difference in estimates at each subsample size compared to the 95%  estimate | | | | | | |

**Table S8: Angolan giraffe - TOD**

| Level | Subsample % | Scaled Mean Degree | | Scaled Mean Strength | | Transitivity | |
| --- | --- | --- | --- | --- | --- | --- | --- |
|  |  | Estimate | Std. Error | Estimate | Std. Error | Estimate | Std. Error |
| Morning | (Intercept) | 0.103 ******* | 0.004 | 0.049 ******* | 0.002 | 0.596 ******* | 0.016 |
|  | 90% | -0.0006 **NS** | 0.006 | -0.0004 **NS** | 0.003 | 0.0004 **NS** | 0.023 |
|  | 70% | -0.0009 **NS** | 0.006 | -0.0004 **NS** | 0.003 | -0.002 **NS** | 0.023 |
|  | 50% | -0.002 **NS** | 0.006 | -0.0007 **NS** | 0.003 | -0.002 **NS** | 0.023 |
|  | 30% | -0.008 **NS** | 0.006 | -0.0037 **NS** | 0.003 | -0.014 **NS** | 0.023 |
|  | 10% | -0.010 **NS** | 0.006 | -0.0048 **NS** | 0.003 | -0.127 ******* | 0.024 |
|  | 5% | -0.022 ******* | 0.006 | -0.0085 ****** | 0.003 | -0.116 ****** | 0.039 |
| Midday | (Intercept) | 0.094 ******* | 0.004 | 0.046 ******* | 0.002 | 0.635 ******* | 0.016 |
|  | 90% | -0.0009 **NS** | 0.005 | -0.0002 **NS** | 0.003 | -0.0004 **NS** | 0.023 |
|  | 70% | -0.0017 **NS** | 0.005 | -0.0008 **NS** | 0.003 | -0.001 **NS** | 0.023 |
|  | 50% | -0.0019 **NS** | 0.005 | -0.0006 **NS** | 0.003 | -0.004 **NS** | 0.023 |
|  | 30% | -0.0047 **NS** | 0.005 | -0.0024 **NS** | 0.003 | -0.028 **NS** | 0.023 |
|  | 10% | -0.015 ****** | 0.005 | -0.0073 ****** | 0.003 | -0.161 ******* | 0.026 |
|  | 5% | -0.015 ****** | 0.005 | -0.0081 ****** | 0.003 | -0.188 ******* | 0.043 |
| Evening | (Intercept) | 0.099 ******* | 0.005 | 0.057 ******* | 0.003 | 0.725 ******* | 0.016 |
|  | 90% | 0.0009 **NS** | 0.006 | 0.0008 **NS** | 0.004 | 0.003 **NS** | 0.023 |
|  | 70% | 0.0003 **NS** | 0.006 | 0.0007 **NS** | 0.004 | 0.0007 **NS** | 0.023 |
|  | 50% | -0.0003 **NS** | 0.006 | -0.0006 **NS** | 0.004 | -0.012 **NS** | 0.023 |
|  | 30% | -0.004 **NS** | 0.006 | -0.0018 **NS** | 0.004 | -0.025 **NS** | 0.023 |
|  | 10% | -0.015 ***** | 0.006 | -0.0073 ***** | 0.004 | -0.149 ******* | 0.026 |
|  | 5% | -0.017 ***** | 0.006 | -0.011 ****** | 0.004 | -0.328 ******* | 0.038 |
|  | Note: P-value: 0.1-1 **NS**, 0.05 *****, 0.01 ******, 0.001 *******  Estimates: Indicate the difference in estimates at each subsample size compared to the 95%  estimate | | | | | | |

**Appendix S5**

**Table S9: Tables of paired t-tests of the fallow deer population – Sampling biased VS random**

| Subsample % | Paired t-test | Metric | | |
| --- | --- | --- | --- | --- |
|  |  | Scaled Mean Degree | Scaled Mean Strength | Transitivity |
| 95% | Mean Difference | - 0.065 | - 0.003 | - 0.029 |
|  | P-value | **< 0.001** ******* | **< 0.001** ******* | **< 0.001** ******* |
|  | T-statistics | - 49.023 | - 10.100 | - 37.251 |
| 90% | Mean Difference | - 0.068 | - 0.004 | - 0.031 |
|  | P-value | **< 0.001** ******* | **< 0.001** ******* | **< 0.001** ******* |
|  | T-statistics | - 51.688 | - 11.307 | - 40.621 |
| 70% | Mean Difference | - 0.067 | - 0.004 | - 0.030 |
|  | P-value | **< 0.001** ******* | **< 0.001** ******* | **< 0.001** ******* |
|  | T-statistics | - 41.130 | - 9.499 | - 31.643 |
| 50% | Mean Difference | - 0.069 | - 0.004 | - 0.032 |
|  | P-value | **< 0.001** ******* | **< 0.001** ******* | **< 0.001** ******* |
|  | T-statistics | - 32.127 | - 7.377 | - 26.126 |
| 30% | Mean Difference | - 0.069 | - 0.004 | - 0.032 |
|  | P-value | **< 0.001** ******* | **< 0.001** ******* | **< 0.001** ******* |
|  | T-statistics | - 20.840 | - 4.717 | - 17.115 |
| 10% | Mean Difference | - 0.064 | - 0.002 | - 0.030 |
|  | P-value | **< 0.001** ******* | **0.035** ***** | **< 0.001** ******* |
|  | T-statistics | - 11.772 | - 2.143 | - 9.985 |
| 5% | Mean Difference | - 0.067 | - 0.003 | - 0.032 |
|  | P-value | **< 0.001** ******* | 0.110 **NS** | **< 0.001** ******* |
|  | T-statistics | - 8.321 | - 1.613 | - 6.381 |
|  | Note: P-value: 0.1-1 **NS**, 0.05 *****, 0.01 ******, 0.001 ******* | | | |
